# Low-cost and clinically applicable copy number profiling using repeat DNA

**DOI:** 10.1101/394429

**Authors:** S Abujudeh, SS Zeki, MCV van Lanschot, M Pusung, JMJ Weaver, X Li, A Noorani, AJ Metz, J Bornschein, L Bower, A Miremadi, RC Fitzgerald, ER Morrissey, AG Lynch, Oesophageal Cancer Clinical and Molecular Stratification (OCCAMS)Consortium, Rebecca C. Fitzgerald, Ayesha Noorani, Paul A.W. Edwards, Nicola Grehan, Barbara Nutzinger, Caitriona Hughes, Elwira Fidziukiewicz, Jan Bornschein, Shona MacRae, Jason Crawte, Alex Northrop, Gianmarco Contino, Xiaodun Li, Rachel de la Rue, Maria O’Donovan, Ahmad Miremadi, Shalini Malhotra, Monika Tripathi, Simon Tavaré, Andy G. Lynch, Matthew Eldridge, Maria Secrier, Lawrence Bower, Ginny Devonshire, Juliane Perner, Sriganesh Jammula, Jim Davies, Charles Crichton, Nick Carroll, Peter Safranek, Andrew Hindmarsh, Vijayendran Sujendran, Stephen J. Hayes, Yeng Ang, Shaun R. Preston, Sarah Oakes, Izhar Bagwan, Vicki Save, Richard J.E. Skipworth, Ted R. Hupp, J. Robert O’Neill, Olga Tucker, Andrew Beggs, Philippe Taniere, Sonia Puig, Timothy J. Underwood, Fergus Noble, Jack Owsley, Hugh Barr, Neil Shepherd, Oliver Old, Jesper Lagergren, James Gossage, Andrew Davies, Fuju Chang, Janine Zylstra, Ula Mahadeva, Vicky Goh, Francesca D. Ciccarelli, Grant Sanders, Richard Berrisford, Catherine Harden, Mike Lewis, Ed Cheong, Bhaskar Kumar, Simon L Parsons, Irshad Soomro, Philip Kaye, John Saunders, Laurence Lovat, Rehan Haidry, Laszlo Igali, Michael Scott, Sharmila Sothi, Sari Suortamo, Suzy Lishman, George B. Hanna, Christopher J. Peters, Anna Grabowska, Richard Turkington

**Author notes:** A full list of contributors from the OCCAMS Consortium is available at the end of the manuscript. These authors contributed equally to this work (S Abujudeh, SS Zeki). These authors contributed equally to this work (RC Fitzgerald, ER Morrissey, AG Lynch).

## Abstract

Large-scale cancer genome studies suggest that tumors are driven by somatic copy number alterations (SCNAs) or single-nucleotide variants (SNVs). Due to the low-cost, the clinical use of genomics assays is biased towards targeted gene panels, which identify SNVs. There is a need for a comparably low-cost and simple assay for high-resolution SCNA profiling. Here we present our method, conliga, which infers SCNA profiles from a low-cost and simple assay.

---

Somatic copy number alterations (SCNAs) are common in cancer. On average, cancer samples see SCNAs in 34% of the genome, with 17% of the genome amplified and 16% deleted [1, 2, 3]. Certain SCNAs, particularly amplifications of oncogenes and deletions of tumor suppressor genes, have been found to be major drivers in tumor development, associated with prognosis and response to therapy [1]. SCNA burden varies considerably between cancer types [3]. For example, oesophageal adenocarcinoma (OAC) has relatively high levels of SCNAs [4, 5, 6, 7], and generally develops from Barrett’s oesophagus. Patients with OAC tend to be diagnosed at a late stage, when spread has occurred to lymph nodes and distant organs. This makes treatment more difficult and leads to poor prognosis [8]. Although most patients with Barrett’s do not progress, early stage disease (high grade displasia or intramucosal adenocarcinoma) can be successfully treated, usually obviating the need for surgery. There is a critical need to develop technologies that can detect early disease and distinguish between patients at low versus high risk for progression. Since most mutations in OAC driver genes are already present in pre-malignent disease [9], but an increased SCNA load distinguishes OAC [10, 11, 12], low-cost SCNA profiling would be a valuable research and clinical tool.

SCNAs have been identified using a number of methods, including comparative genomic hybridisation (CGH) [13], array-based CGH [14], single nucleotide polymorphism (SNP) arrays [15], and whole-genome sequencing (WGS) [16]. Recently, low-coverage (LC) WGS has gained popularity due to its reduced cost and strong performance [17]. However, while LC WGS reduces the cost of sequencing, standard WGS library preparation is required with its associated fixed expense and time needed to produce each sample. A technically simple, fast, easily automated, high-resolution and inexpensive alternative method for SCNA detection, with clinical potential, would be extremely valuable.

Recent studies have shown the genome can be amplified at multiple (>10,000) genomic loci with the use of a single non-specific primer pair, using the FAST-SeqS method [18, 19]. With this approach, two polymerase chain reaction (PCR) rounds replace the complicated and expensive library preparation steps associated with WGS. The amplified regions are sufficiently short such that the assay can be performed on cell-free DNA as well as DNA extracted from tissue biopsies. The resulting amplicons can be sequenced, with samples multiplexed on the same sequencing lane. With this method, we maintain a similar sequencing depth to 30-50X high-coverage (HC) WGS while sequencing only specific loci. This is in contrast to LC WGS which samples the whole genome but at reduced sequencing depth (Supplementary Fig. 1). The cost involved in sample preparation and sequencing combined is approximately £14 per sample compared with approximately £52-72 for LC WGS, depending on the library preparation kit used (Supplementary Note 1). The sample preparation can be performed in less than an hour with minimal hands-on time, compared to approximately 3 hours or greater for LC WGS.

Until now, the use of FAST-SeqS data has been limited to the detection of whole chromosome gains [18] and entire chromosome arm gains and losses [19, 20]. This means that chromosome segment (focal) alterations are not detected, or perhaps falsely considered as whole chromosome or chromosome arm alterations. Moreover, in these methods SCNAs are not quantified and regions are simply classified as amplified, deleted or normal.

Here we present a method (and associated tool: ‘conliga’) that uses a fully probabilistic approach to infer relative copy number (RCN) alterations at each locus from FAST-SeqS data. conliga provides a RCN profile per sample and therefore enables this low-cost sequencing approach to be used as a SCNA assay.

Based on observations of raw data (Supplementary Note 2, Supplementary Fig. 1), we created a probabilistic model (Methods, Supplementary Note 3). The model takes account of the observed bias in loci counts, which predominantly results from unequal PCR efficiencies between loci. Since neighboring loci are likely to share the same copy number, we use a hidden Markov model (HMM) to model the spatial dependence between loci. This allows loci with high counts to share statistical strength with neighboring loci, enabling us to infer contiguous regions of copy number more accurately. Moreover, we use a Bayesian nonparametric approach (sticky HDP-HMM) [21] to address the issue of the unknown number of copy number levels present in a given sample a priori (Methods). We use Markov Chain Monte Carlo (MCMC) methods to infer the RCN of each locus, plus all other latent variables in the model (Methods, Supplementary Table 1, Supplementary Notes 4, 5 and 6). This enables us to provide the uncertainty of the RCN estimates, summarized by credible intervals, in conliga’s standard output.

To test our method, we analysed 11 oesophageal adenocarcinoma tumors (Methods, Supplementary Tables 2 and 3), which had been sequenced using HC WGS (>50X) and FAST-SeqS. In addition, we downsampled the WGS data of each sample to nine million reads to simulate typical LC WGS (∼ 0.1X coverage) samples (Methods). We compared the copy number calls derived from ASCAT [22] (applied to HC WGS data) with the RCN calls from QDNAseq [17] (LC WGS data) and conliga (FAST-SeqS data). conliga and QDNAseq achieved a median Pearson correlation coefficient with ASCAT of 0.95 and 0.98 respectively (Methods, Supplementary Table 4).

In figure 1a-d we demonstrate that similar RCN profiles are obtained with the three methods for an example sample (OAC2) and that high-resolution SCNA information is maintained by sampling genomic loci using FAST-SeqS. Figure 1e and 1f show the performance of conliga and QDNAseq, both obtaining similar Pearson correlation coefficients with ASCAT’s RCN calls across all 11 OAC samples (conliga: 0.953, QDNAseq: 0.987) and residual distributions when compared to ASCAT (Methods). It should be noted that by downsampling reads from the same WGS sample, this analysis is potentially biased in favor of QDNAseq’s results.

**Figure 1.**
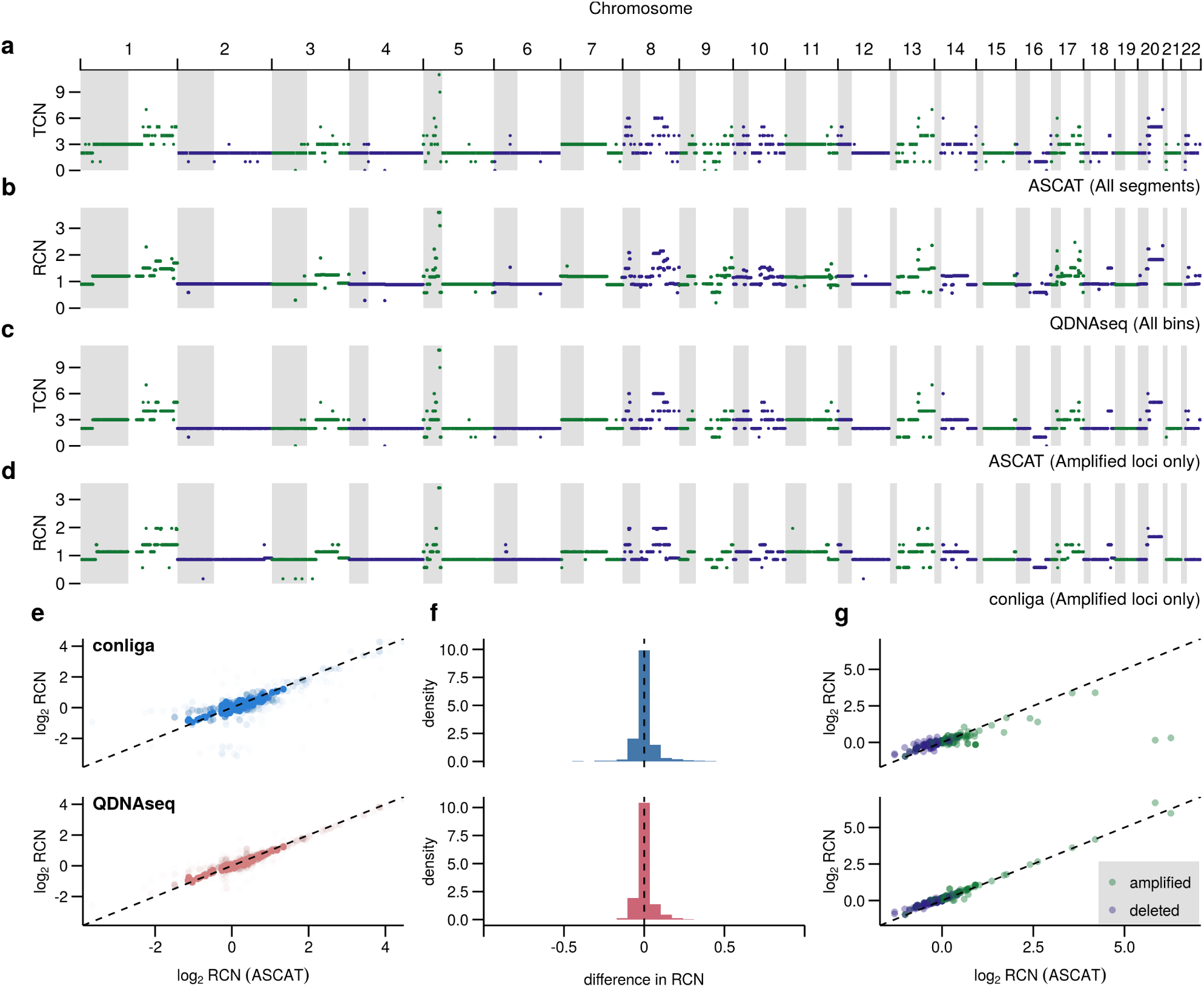
Comparison of conliga method with ASCAT and QDNAseq. (a) Total copy number profile determined by ASCAT from HC WGS data for sample OAC2, showing all copy number segments. (b) Relative copy number profile determined by QDNAseq from LC WGS data for sample OAC2, showing all 15 Kbp bins. (c) Total copy number profile determined by ASCAT from HC WGS data for sample OAC2, showing ASCAT’s copy number calls at the intersection of ASCAT’s called regions and FAST-SeqS loci. (d) Relative copy number profile determined by conliga from FAST-SeqS data for sample OAC2, at the intersection of ASCAT’s called regions and FAST-SeqS loci. (e) Comparison of log_2_ relative copy number calls from 11 samples between conliga and ASCAT (top) and QDNAseq and ASCAT (bottom). All RCN calls at the intersection of ASCAT’s called regions, QDNAseq 15Kb bins and FAST-SeqS loci in all 11 OAC samples are shown as points. (f) Distribution of differences between ASCAT RCN calls and conliga RCN estimates for 11 OAC samples (top) and ASCAT RCN calls and QDNAseq RCN estimates for 11 OAC samples (bottom). (g) Comparison of performance at gene level resolution between ASCAT and conliga (top) and ASCAT and QDNAseq (bottom). The values represent the weighted mean of RCN calls at each gene for each of the 11 OAC samples (Methods).

From the literature [23, 12] we selected a set of 36 genes that have been observed to be recurrently amplified or deleted in OAC (Supplementary Table 5, Methods). We determined the weighted mean of the RCN calls for these genes for each sample via each method (Methods, Supplementary Tables 6 and 7). While FAST-SeqS/conliga would not be the assay of choice if only interested in a small gene panel, in Figure 1g we see that there are only two instances from 396 comparisons (36 genes x 11 samples) where a substantially different result would be achieved. Naturally if an SCNA is so narrow as to fall between two FAST-SeqS loci then it will not be detected in this way, but the detection of many highly-localized events demonstrates how informative FAST-SeqS/conliga can be. Even within this panel of 36, it is notable that some genes harbour FAST-SeqS loci (Supplementary Tables 8 and 9), providing evidence of intra gene SCNAs in some cases, such as the focal deletions observed in FHIT, PARK2, and MACROD2 (Supplementary Fig. 2). Focal deletions such as these may be functionally relevant, potentially rendering tumor suppressor genes inactive.

The purity of tumor samples obtained by dissection can vary widely [24], as can samples obtained non-invasively, e.g ctDNA from plasma [25]. As tumor purity reduces, the copy number signal to noise ratio decreases. To determine the performance of conliga and QDNAseq under different purity conditions, we generated samples with varying purity by mixing sequencing reads from normal and OAC samples (Methods). FAST-SeqS samples were generated with two million reads and LC WGS samples were generated with nine million reads.

Figure 2a shows the performance of both methods for sample OAC3. At 30% purity, both conliga and QDNAseq recapitulate the copy number profile as determined by ASCAT. At 5%, other than the focal amplification on chromosome 12, QDNAseq fails to detect sub chromosomal SCNAs, whereas conliga shows evidence of chromosome arm and sub-chromosomal arm changes. At 2% purity, conliga is able to distinguish some of the more prominent chromosomal arm SCNAs. The focal amplification on chromosome 12 is identified by conliga at 0.75% and 0.5% purity and not detected by QDNAseq below 1%. At 0.75%, 0.5% and 0% purity, it is hard to distinguish whole chromosome SCNAs from noise generated by segmentation in the QDNAseq profiles. This highlights the advantage of conliga’s ability to assign loci to discrete states, meaning we can easily distinguish when SCNAs are and are not different between loci. Despite using 4.5 fold fewer reads, conliga appears to be more sensitive than QDNAseq.

**Figure 2:**
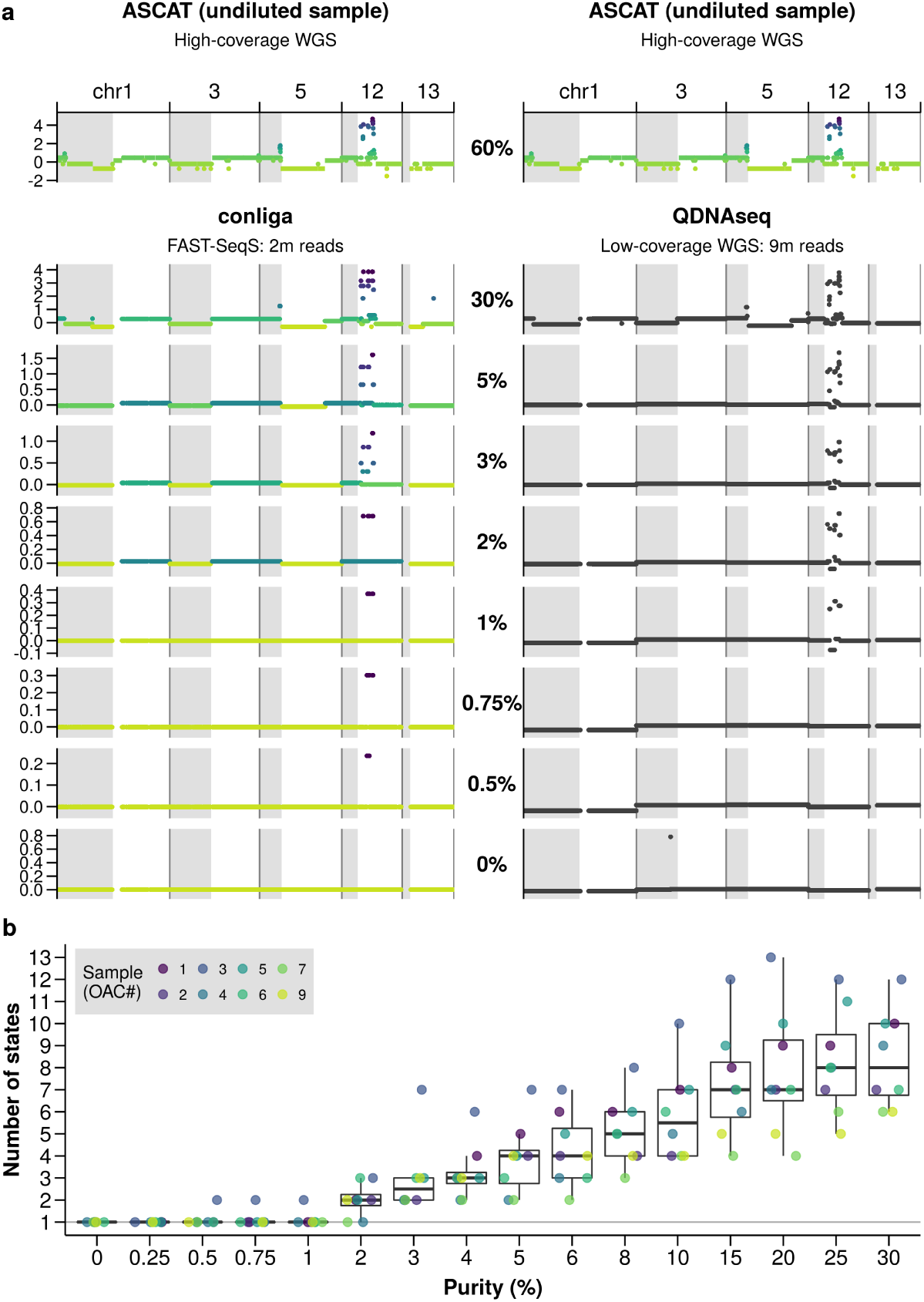
Comparing the performance of SCNA detection in low tumor purity samples and determining the limit of detection. (a) left column: relative copy number calls by conliga at different dilutions of sample OAC3, compared to ASCAT relative copy number profile (top left), discrete copy number states are colored with a gradient (light green to purple), highlighting regions with differing SCNAs. right column: relative copy number calls by QDNAseq at different dilutions of sample OAC3, compared to ASCAT relative copy number profile (top right). (b) The number of copy number states detected by conliga in each of eight OAC samples at differing purity levels. The limit of detection is determined by the lowest purity level in which more than one copy number state is detected.

In Figure 2b, we show that conliga is able to detect SCNAs at 3% purity in all samples (eight), five at 2% and one at 0.5%. The limit of detection is dependent on the amplitude and lengths of SCNAs present in the sample. Long chromosomal arm amplifications can be detected at 2-3% purity, while some focal amplifications (particularly those occurring at loci with a bias towards obtaining a high number of counts) can be detected at <1% purity (e.g. chr12 in OAC3, Figure 2a). The limit of detection also depends on the technical variability of the protocol and the total number of reads per sample. Increasing the total number of reads beyond two million and reducing technical variability would further improve the limit of detection.

These data demonstrate the potential clinical utility of FAST-SeqS coupled with conliga. Ciriello et al. identified that either somatic single nucleotide variants (SNVs) or SCNAs [3] can drive oncogenesis. Currently, there is a bias towards screening for SNVs using targeted gene panels [26] meaning SCNA-driven cancers may not be detected. To this end, we analyzed samples with pre-malignant disease (Barrett’s oesophagus) and were able to detect clinically relevant copy number alterations, such as evidence for focal gains of PRKCI, ERBB2 and GATA6 and deletions of regions containing CDKN2A, PTPRD, SMAD4 and TP53 (Supplementary Fig. 2). This suggests that there is potential for FAST-SeqS to be used alongside existing low-cost gene panels to detect SCNAs, in addition to SNVs, to screen and surveil patients for the development of cancer.

In addition to use as a detection tool, inexpensive production of FAST-SeqS data allows for large cohorts of patients to be studied to find relationships between SCNA profiles and response to therapies, for example. With this in mind, we looked at the average SCNA profiles across small cohorts of patients with OAC, Gastric cancer and Barrett’s oesophagus (Supplementary Fig. 2, Methods) which highlighted amplifications of known oncogenes such as EGFR, MYC, GATA4, and MDM2, some with known drug targets, and deletions of tumor suppressor genes, e.g. FHIT, TP53, SMAD4 and RUNX1. Other potential uses include low-cost screening of samples in large-scale cancer genomes studies, such as ICGC or TCGA projects, prior to further genomic analyses. Furthermore, due to the low-cost and low-input DNA required, several spatially or temporally related samples can be analyzed for the purposes of determining how SCNAs accumulate in normal tissues and contribute to tumor evolution, in a similar fashion to previous studies on somatic mutations in the eyelid epidermis [27].

Areas for future study could include determining an acceptable number of reads which balances the cost and limit of detection, finding ways to minimise the technical variability, and altering the number of reads obtained at specific loci to increase statistical power in regions of interest.

We have shown that FAST-SeqS data can be used as a viable, inexpensive, and simple alternative to LC WGS for the purpose of SCNA detection and quantification. conliga provides accurate and high-resolution SCNA profiles across the genome and at regions of interest such as oncogenes and tumor suppressors. conliga (applied to FAST-SeqS data with two million reads per sample) is particularly useful in detecting and discriminating SCNAs in low purity samples and our results suggest it to be more sensitive than QDNAseq (using LC WGS, nine million reads) for this purpose. We believe that conliga makes FAST-SeqS data a clinically valuable diagnostic assay to detect and monitor patients for the development of cancer, as well as a useful research tool, enabling inexpensive and fast SCNA profiling of cancer samples.

## Methods

### conliga: statistical model

#### Statistical model for sample counts

We model the sample counts, in *L* selected loci, by assuming that the count at locus *l* in chromosome arm *r* in sample *j* is distributed:

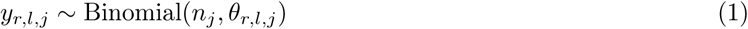

Here, *n*_*j*_ is the total number of sequencing reads aligned to the *L* loci in sample *j, θ*_*r,l,j*_ represents the probability of observing an aligned read at locus *l* in chromosome arm *r* in sample *j*. We model *θ*_*r,l,j*_ as follows:

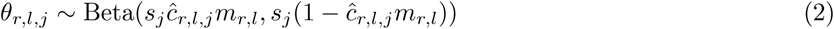

Here, *s*_*j*_ is the inverse dispersion variable for sample *j* where *s*_*j*_ > 0, *m*_*r,l*_ represents the probability of an aligned sequencing read originating from locus *l* in chromosome arm *r* in a control sample, where 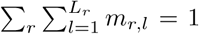 and ĉ_*r,l,j*_ is the relative copy number at locus *l* in chromosome arm *r* in sample *j*. The number of loci in each chromosome arm is denoted as *L*_*r*_ and so the total number of loci, *L* = Σ*_r_ L*_*r*_.

We can interpret *m* as defining the bias in observing aligned read counts from the FAST-SeqS protocol. This bias can be explained by unequal PCR efficiencies between loci in addition to biases in aligning reads uniquely to FAST-SeqS loci, among other factors. Note that:

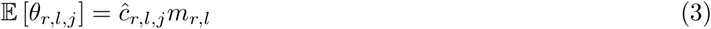

We can be interpret this equation intuitively; the relative copy number scales the probability of reads to align to a locus. For example, if the relative copy number of a locus is 2 we expect the proportion of reads at the locus to double. This fits with our observations shown in Supplementary Fig. 1.

The inverse dispersion variable, *s*_*j*_, is sample specific and reflects our observations that the level of dispersion varies between samples. This variation in dispersion between samples might be due to varying levels of DNA degradation and/or varying quantities of starting material between samples, among other factors. *s*_*j*_ relates to the variance and the mean of *θ*_*r,l,j*_ in the following way:

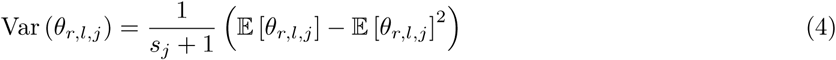

The expected count, *y*_*r,l,j*_, in chromosome arm *r* at locus *l* in sample *j* is:

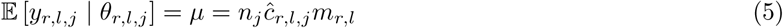

The variance of *y*_*r,l,j*_ can be written as a quadratic function of *µ* with the coefficients being a function of *n*_*j*_ and *s*_*j*_:

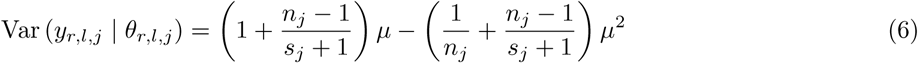

Note that in the limit *s*_*j*_ *→* ∞, a Binomial noise model is recovered.

#### Probabilistic generative model of loci counts for control samples

We assume that the loci within a control sample, *k*, have equal copy numbers (diploid). This means that the RCN for each locus is 1. By setting *ĉ*_*r,l,k*_ = 1, we model the generative process of counts from a control sample as follows:

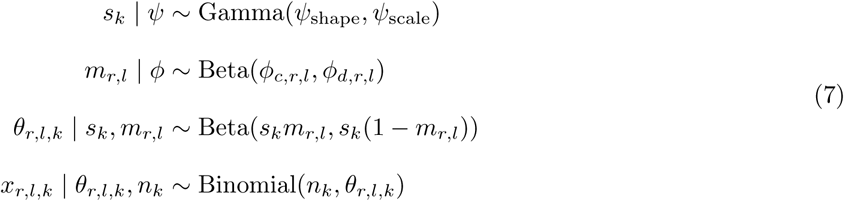

Here, Gamma(*ψ*_*shape*_, *ψ*_*scale*_) represents the prior distribution over the sample specific inverse dispersion parameter, *s*_*k*_, and Beta(*ϕ*_*c,r,l*_, *ϕ*_*d,r,l*_) defines the prior distribution over *m*_*r,l*_.

#### Linking FAST-SeqS loci using a hidden Markov model

We assume that chromosome arms are independent. By that we mean, the RCN of the first locus in arm q is independent of the RCN of the last locus in arm p from the same chromosome (and all other chromosome arms). As such, we model each chromosome arm as an independent Markov chain for each sample *j*. We denote (note that for simplicity we have dropped the sample index *j*):

- *z*_*r,l*_ as the *hidden state* (or *copy number state*) of the Markov chain at locus *l* in chromosome arm *r*
- *π*^0^ as the *initial distribution* of the first locus (*l* = 1), in chromosome *r*
- *π*_*u*_ as the *transition distribution* for hidden state, *u*
- *ĉ*_*u*_ as the *relative copy number* associated with hidden state, *u*.

The first locus of a chromosome arm (*l* = 1) is distributed:

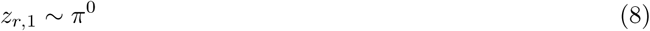

For all other loci (*l* > 1):

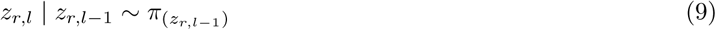

The count, *y*_*r,l*_, at locus *l* in chromosome arm *r* is conditionally independent of the hidden states and observations of other loci:

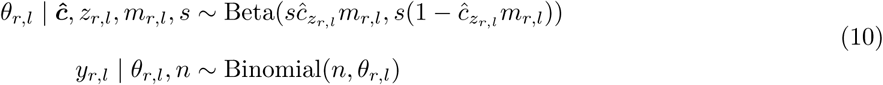

The joint density for *L*_*r*_ loci in chromosome arm *r* is:

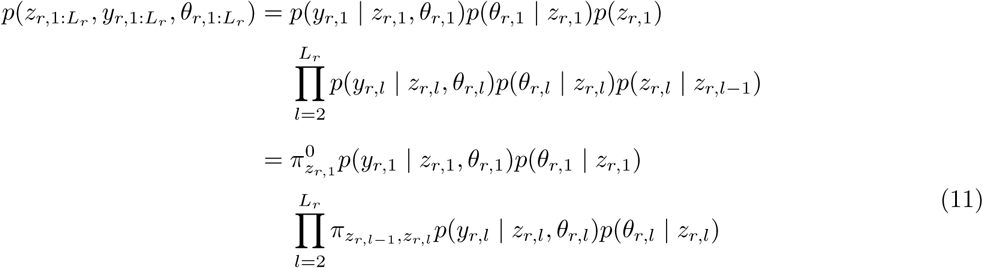

where, *z*_*r,*1:*L_r_*_ denotes the sequence {*z*_*r,*1_, *…, z*_*r,L_r_*_ }, *y*_*r,*1:*L_r_*_ denotes {*y*_*r,*1_, *…, y*_*r,L_r_*_ }, and *θ*_*r,*1:*L_r_*_ denotes {*θ*_*r,*1_, *…, θ*_*r,L_r_*_ }. The joint density for all *L* loci in the genome is given by:

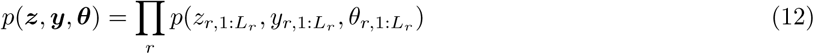

#### Probabilistic generative model of a sample’s relative copy number profile

The number of copy number states present in a sample is unknown a priori. In samples that have equal copies of each locus, only one copy number state is present. Conversely, it is possible (although unlikely) that each locus has its own unique copy number, meaning that there could be up to *L* copy number states in a sample. Additionally, we expect neighboring loci to share the same copy number given their genomic distance from each other (Supplementary Fig. 1). To address these two features of the data, we used the sticky hierarchical Dirichlet process hidden Markov model (sticky HDP-HMM) [21] as a framework to model the generative process of a sample’s relative copy number profile. By doing so, we adequately model the spatial persistence of copy number states and allow for countably infinite numbers of states within a sample. The generative model is as follows:

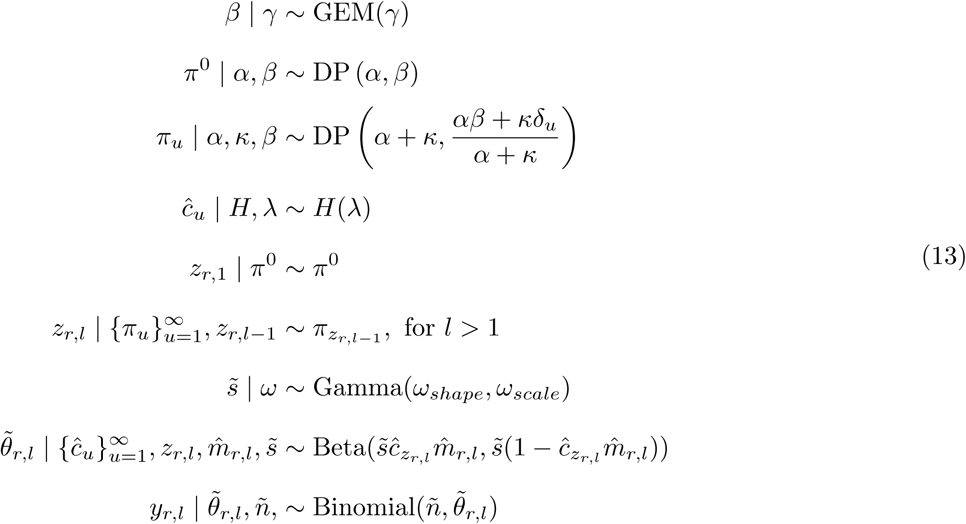

Note that we use 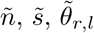 to distinguish these variables from those in the probabilistic model of control counts (equation 7) and denote them as specific to the sample with copy number profile. Here, GEM denotes the stick-breaking construction of the Dirichlet Process as described in Fox *et al.* [21]. *γ* is a hyperparameter of the sticky HDP-HMM and represents our prior on the number of copy number states in the sample; the greater the value of *γ*, the greater number of copy number states we expect in the sample. Each row of the transition matrix, *π*_*u*_, is drawn from a Dirichlet Process and depends on *β, α* and *κ*. It can be shown that:

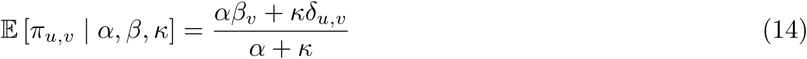

where *δ*_*u,v*_ represents the discrete Kronecker delta function. If we define 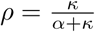 (as in Fox *et al.* [21]) and by noting that *α* = (1 − *ρ*)(*α* + *κ*), we obtain:

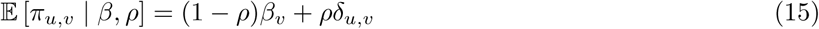

As such, we see that *ρ* defines how much weight is placed on self-transition within a copy number state. The vector, *β*, itself drawn from a Dirichlet Process, represents the global transition distribution and holds information about the proportion of loci expected in each copy number state.

The variance of the transition probability from copy number state *u* to *v* is given by:

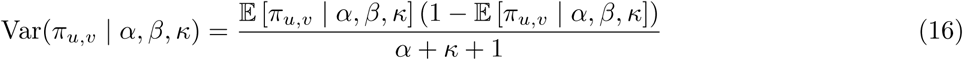

We see that *α* + *κ* is inversely proportional to the variance of the state transition probabilities.

*H* is the prior base distribution of the Dirichlet Process and represents a parametric distribution, which in this case is a Gamma distribution, with parameters *λ*. It can be viewed as our prior probability distribution on the relative copy number values of the hidden states.

Note that 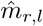 refers to the maximum a posteriori (MAP) value of *m*_*r,l*_ and is such assumed to be a known quantity in equation 13. For simplicity, the hyperparameters (*α, κ, γ, λ, ω* and *n*) are shown as fixed quantities in the model. In practice, *γ, λ, ω* and *n* are treated as fixed, while the model is parameterized in terms of *ρ* and (*α* + *κ*), with a Beta prior placed on *ρ* and a Gamma prior placed on (*α* + *κ*) as in Fox *et al.* [21]. See the section on inference for further details of prior distributions used and Supplementary Note 3 for further discussion on the model.

### Inference

#### Inference of loci count proportion bias (*m*)

Given a set of *K* control samples, and their loci counts, *x*_*k*_, we used our model defined in equation 7 and Markov Chain Monte Carlo (MCMC) methods to infer the latent variables *m* and *s* (the vector of sample specific inverse dispersion parameters). A Metropolis-Hastings MCMC algorithm was used to obtain a sample of the posterior probability of *m*_*r,l*_ for all r and l, and *s*_*k*_, for each sample *k*. Full details of the algorithms are provided in Supplementary Notes 4 and 5. Count data for samples analyzed in this study, processed by the pipeline described, are provided in Supplementary Table 10.

For each sequencing experiment, a suitable set of controls samples were used (see Supplementary Table 11 for the list of samples used in each experiment). As described in equation 7, control samples were assumed to have a relative copy number of one at each locus. In all experiments described in this paper, we used the following values for the hyperparameters:

- *ψ*_*shape*_ = 1.5, *ψ*_*scale*_ = 10^6^; where *ψ*_*shape*_ and *ψ*_*scale*_ define the shape and scale of the Gamma prior distribution on *s*_*k*_, respectively.
- *ϕ*_*c,r,l*_ = 1 and *ϕ*_*d,r,l*_ = 1 for all *r* and *l*; i.e. we used a flat Beta(1, 1) prior for all *m*_*r,l*_

In each sequencing experiment, 20,000 iterations of the MCMC were run and the first 5,000 iterations were discarded (burn-in). Maximum a posteriori (MAP) estimates of *m* (denoted as 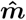) were obtained by determining the mode of the sampled posterior densities for each locus using the KernSmooth R package [28]. Note that the MAP estimates are unlikely to sum to 1 exactly, and as such we enforced this by setting 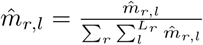.

#### Inference of relative copy number profile

Given 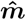 and the loci counts (*y*) for a sample with unknown copy number profile, we used the generative model defined in equation 13 and MCMC methods (based on algorithm 3 in Fox *et al.* [21]) to infer the latent variables in our model. MCMC methods were used to obtain a sample of the posterior probability of the hidden state of each locus (*z*_*r,l*_ for all *r* and *l*), the relative copy number of each hidden state (*ĉ*_*u*_), the sample specific inverse dispersion 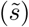, along with other latent variables in our generative model. Full details of the MCMC algorithms can be found in Supplementary Notes 4 and 6. In all experiments described in this paper, we used the following values for the hyperparameters:

- *γ* = 1
- Gamma(2000, 10) prior distribution (defined by shape and scale) was placed on (*α* + *κ*)
- Beta(100000, 100) prior was placed on *ρ*
- Gamma(3, 1) prior distribution (defined by shape and scale) was placed on the relative copy number value of the hidden states; the shape and scale parameters are defined by *λ* in equation 13
- *ω*_*shape*_ = 1.5, *ω*_*scale*_ = 10^6^; where *ω*_*shape*_ and *ω*_*scale*_ define the shape and scale of the Gamma prior distribution on 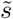, respectively

The output of the MCMC was summarized in two main ways, 1) by marginalizing out the copy number state information and computing the MAP estimate (using KernSmooth R package [28]) and credible interval of the relative copy number of each locus, 2) by making use of the copy number state assignments in the following way:

1. we determined the MAP number of states observed in the MCMC chain (after burn-in). This was achieved by calculating the number of populated states in each iteration of the MCMC, and then choosing the most frequently observed number of populated states. Note that a state was considered populated in an iteration of the MCMC if at least one locus was assigned to it.
2. we filtered the iterations of the MCMC (after burn-in), choosing only those iterations that had the number of populated states equal to the MAP number of states.
3. we used the Stephens algorithm (algorithm 2 in the paper) [29] along with the Hungarian (Munkres) algorithm [30] to relabel the states, to resolve the label switching problem inherent in MCMC methods.
4. we calculated the MAP estimate and credible intervals for the relative copy number values of each relabeled state.
5. we assigned each locus to a relabeled state, choosing the relabeled state it was most frequently assigned to in the filtered iterations of the MCMC chain.

For the results presented in Figure 2, summarization method 2 was used. For all other results presented in the paper, summarization method 1 was used. For the oesophageal cancer, gastric cancer and Barrett’s oesophagus samples, 50,000 iterations of the MCMC were run and the chain was thinned such that every 5th iteration of the MCMC was output to file. Additionally, the first 20,000 iterations of the MCMC were discarded (burn-in), to ensure the Markov chain had reached its equilibrium distribution. For the in silico diluted samples, presented in Figure 2, 30,000 iterations were run, with the chain thinned so that every 5th sample was output to file and the first 5,000 iterations of the MCMC were discarded.

### Sample preparation and sequencing of samples

#### Sample preparation and generation of FAST-SeqS data

Sequencing libraries were prepared using two rounds of PCR, using a similar protocol to previously published methods [18, 19]. Each extracted DNA sample underwent a 50 µl first round PCR reaction with 10 µl 5x Phusion HF Buffer (ThermoFisher Scientific), 1 µl 10 mm dNTP (ThermoFisher Scientific), 5 µl of both the forward and reverse primers (0.5 µm) each (Sigma-Aldrich), 0.5 µl Phusion Hot Start II DNA Polymerase 2U/µl, 5-10 µl DNA template depending on the extracted concentration, and RNAse free water to make the total reaction volume. The cycling conditions for the L1PA7 primers were 98 °C for 120 s followed 2 cycles of 98 °C for 10 s, 57 °C for 120 s, and 72 °C for 120 s. The second round was also carried out as a 50 µl sample reaction using 20 µl taken from the first round. The rest of the reaction constituents were the same as the first round reaction with the exception of primers (Supplementary Table 12), which contained a unique index for each sample. The cycling conditions for the second round reaction were 98 °C for 120 s followed by 13 cycles of 98 °C for 10 s, 65 °C for 15 s, and 72 °C for 15 s for all the primers. After the second round, samples underwent quantification using the 2200 TapeStation (Agilent), Agilent 2100 Bioanalyser (Agilent) and Kapa quantification (KapaBiosystems) prior to submission for sequencing. The samples were then pooled in equimolar concentrations and gel extracted according to manufacturer’s instructions (Qiaquick gel extraction kit, Qiagen). Finally the samples were submitted for sequencing on a MiSeq (Illumina) platform. All samples were run with 20% PhiX to increase complexity for sequencing. Sequencing was performed as 150bp single end. Samples were run with at least three normal controls prepared at the same time and sequenced on the same platform.

#### Sample preparation and generation of high-coverage WGS data

WGS library preparation and sequencing was performed as previously described by Secrier *et al.* [6].

#### In silico generation of low-coverage WGS data

For our purposes, LC WGS data was defined as nine million single-end 50 base pair reads per sample because this was the type of data analyzed in Scheinin *et al.* [17]. Samples are typically multiplexed together and sequenced on a single Illumina sequencing lane. After processing and alignment of the reads, we expect approximately 0.1X coverage of the genome (as per analysis described in Scheinin *et al.*). We obtained LC WGS data by down-sampling reads from HC WGS BAM files in the following way:

1. we selected a subset of the alignments, containing only reads sequenced on a single lane (chosen to be the lane from the first read in the BAM file), and trimmed the reads and Phred scores to the first 50 base pairs using a custom Bash script.
2. The resulting alignments were filtered (using samtools [31] version 0.1.18), excluding those that were secondary alignments (-F 256) and including only those that were first in a pair (-f 64) and output to a new BAM file.
3. This BAM file was down-sampled to 9 million reads/alignments using the DownsampleSam command from Picard tools (http://broadinstitute.github.io/picard, version 2.9.1) using the “Chained” strategy.
4. The resulting BAM file was converted to FASTQ by SamToFastq (Picard tools).
5. The FASTQ file was aligned to GRCh38 (GenBank accession: GCA_000001405.15, no alt analysis set) using BWA-backtrack (bwa samse and bwa aln, version 0.7.15-r1140) [32], which is more suitable for reads below 70 base pairs in length.
6. In the resulting BAM file, we removed PCR duplicates and removed alignments with mapping quality below 37 as per the analysis undertaken by Scheinin *et al.* [17] using samtools (version 0.1.18).

We performed these steps for 11 oesophageal samples and their matched normal samples along with an additional four normal samples obtained from other patients (Supplementary Table 1). This resulted in greater than seven million primary alignments per sample.

#### In silico generation of FAST-SeqS dilution data

We performed an in silico dilution of FAST-SeqS data by mixing sequencing reads from control samples with reads from OAC samples. Since the number of reads in the matched controls were insufficient to create samples with two million reads, we created a pool of control reads (in silico) which were used to dilute the OAC samples. This was done by sub-sampling two million reads from 12 control samples (which were prepared and sequenced in the same batch as the OAC samples). The total number of reads from these 12 control samples was 14,405,596. To obtain a pool of 2 million reads, we used the ‘sample’ command from seqtk (urlhttps://github.com/lh3/seqtk, version: 1.2-r101) to sample a proportion (2*/*14.405596) of each control sample’s reads and merged these together into a single FASTQ file. The reads that were sub-sampled were removed from the control samples (using a custom python script) to avoid using the same reads to fit *m*.

We mixed the pool of control reads with the OAC samples in varying proportions to achieve a desired diluted tumor purity. The OAC samples did not have a tumor purity of 100%, instead they were themselves a mixture of tumor and normal DNA. The purity of these samples were determined by ASCAT-NGS (version 2.1) [22]. Based on ASCAT’s purity value, we calculated the number of reads required from the OAC sample to achieve a desired dilution and total number of reads. This was calculated as follows:

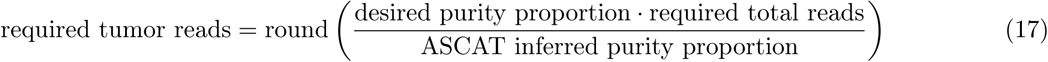

Hence, the number of control reads required were:

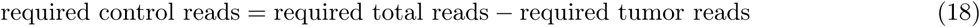

We produced in silico dilution FASTQ files in the following way:

1. we used the ‘sample’ command from seqtk to sample the required number of tumor reads from the OAC FAST-SeqS FASTQ file
2. we used the ‘sample’ command from seqtk to sample the required number of control reads from the pooled control reads FASTQ file
3. We merged the sampled tumor and control reads into a single FASTQ file

We performed these steps for each OAC sample to create diluted samples with two million total reads and the following purity values: 0.3, 0.25, 0.2, 0.15, 0.1, 0.08, 0.06, 0.05, 0.04, 0.03, 0.02, 0.01, 0.0075, 0.005, 0.0025 and 0. Here purity is defined as the proportion of tumor reads in the sample. Of the 11 OAC samples, 8 (OAC1-7 and 9, Supplementary Table 1) were of sufficient initial tumor purity to feasibly create all the desired dilution levels.

#### In silico generation of LC WGS dilution data

We produced in silico diluted LC WGS tumor samples by mixing reads from tumor and matched normal LC WGS BAM files (previously downsampled and filtered as described above). We first calculated the number of reads in the tumor BAM and normal BAM files using samtools (`samtools view -F 256 -c [BAM file]`). Next, we calculated the number of reads required using equations 17 and 18. Using the DownsampleSAM command (Picard tools) and the ‘HighAccuracy’ strategy, we sampled the corresponding desired proportion of reads from the tumor BAM file and normal BAM file. We used samtools to merge the resulting sampled tumor BAM file with the normal BAM file into a single file representing the diluted sample. We aimed to obtain seven million filtered primary alignments per diluted sample (as this is what we expect from nine million reads after alignment and filtering) and dilution levels which matched the diluted FAST-SeqS samples. This was performed for 8 OAC samples and their matched normals (OAC1-7 and 9).

### Processing of FAST-SeqS sequencing data to counts

Each sequencing run of the Illumina MiSeq platform produced a BCL file which was converted to FASTQ format (using Illumina’s bcl2fastq tool). Sequencing reads that failed the Illumina chastity filter were removed. The FASTQ file was demultiplexed into separate FASTQ files corresponding to each sample using the demuxFQ tool (https://genomicsequencing.cruk.cam.ac.uk/glsstatic/lablink/downloads/DemultiplexingGuide.html) with the default settings. The sample barcodes are provided in Supplementary Table 12. Each sample’s FASTQ file was then processed through a custom pipeline which we describe below.

#### Identifying forward primer position

For each read in the FASTQ file, the position of the forward primer sequence was detected by searching for the sequence with the minimum hamming distance to the forward primer sequence using a sliding window. Reads with a minimum hamming distance greater than 5 were discarded.

#### Read trimming

The portion of the reads before and including the forward primer sequence were trimmed. The ends of the reads were also trimmed such that the length of the reads used for downstream analyses were 100 base pairs minus the forward primer length. Any reads shorter than 100 base pairs minus the forward primer length after trimming were discarded.

#### Quality control

After trimming, reads were discarded if they contained at least one base with a Phred quality score less than 20 and/or contained one or more ambiguous base calls (N).

#### Obtaining unique sequences and counts per unique sequence

To avoid aligning the same sequence multiple times, only unique read sequences were kept. For each unique read, the number of identical fragments were recorded.

#### Alignment of unique sequences

Unique raw read sequences were aligned with Bowtie 1.0.0 [33] (using the option: -r). Three mismatches were permitted (option: -v3) and reads aligning to multiple locations were discarded (option: -m1). The reads were aligned to GRCh38 (GenBank accession: GCA_000001405.15, no alt analysis set).

#### Counts and alignments combined

Each sample’s unique read alignments and their corresponding unique read counts were combined into a single file consisting of a matrix of counts. The rows corresponded to genomic positions (the union of genomic positions from the alignments in all samples) and columns corresponded to samples. The first three columns of the matrix corresponded to the chromosome, position and strand for the locus, respectively. The matrix of counts used in this analysis can be found in the conliga R package and in Supplementary Table 10.

### Selecting loci

Rows of the count matrix corresponding to genomic loci within chromosomes X, Y and within unplaced or unresolved contigs were discarded. For each batch of samples, genomic loci obtaining a zero count in any one of a set of control samples were also discarded. Depending on the sequencing batch we analyzed and the controls chosen to filter loci (Supplementary Table 11), this resulted in approximately 10,000 - 12,000 genomic loci across chromosomes 1 to 22.

### Analysis of copy number from FAST-SeqS data

conliga (version 0.1.0) was used to analyze all FAST-SeqS samples in this study (Supplementary Table 1) using R (version 3.2.3) [34] and RcppAramdillo (version 0.6.500.4.0) [35]. Of the 15 OAC samples sequenced, four were excluded due to their obtaining fewer than 350,000 reads. Two control samples were excluded due to their inferred RCN profiles having two main hidden states incompatible with their supposed ‘normal’ status. The values for the priors used and MCMC settings are stated in the inference sections above. The samples used as a basis to filter loci and fit 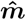 for each experiment are listed in Supplementary Table 9.

### Analysis of copy number from high coverage WGS data

High coverage WGS samples were processed and aligned using BWA-MEM [36] (version 0.5.9) and total copy number (TCN) profiles and normal contamination estimates were provided by ASCAT-NGS (version 2.1) using a pipeline previously described by Secrier *et al.* [6]. The only exception to this was that the reads were aligned to GRCh38 (GenBank accession: GCA_000001405.15, no alt analysis set) rather than GRCh37.

### Analysis of copy number from low-coverage WGS data

QDNAseq (version 1.6.1) was used to obtain relative copy number calls for all LC WGS data. The bin size used was 15Kb as per the analysis performed in Scheinin *et al.* [17] for 0.1X LC WGS. The bins were created using GRCh38 (BSgenome.Hsapiens.NCBI.GRCh38) and a mappability file (bigWig format) for 50-mers was created for GRCh38 using the GEM library (GEM-binaries-Linux-x86_64-core_i3-20130406-045632) https://sourceforge.net/projects/gemlibrary/. 15 normal LC WGS samples (Supplementary Table 1), were used to run the applyFilters and iterateResiduals functions. 11 of these 15 samples correspond to the matched normals of the oesophageal samples (Supplementary Table 1). We did not run the functions normalizeBins and normalizeSegmentedBins which scale the read counts by the median value. This was not necessary and would make the comparison between ASCAT, QDNAseq and conliga results more difficult to interpret.

### Comparison of copy number between methods

ASCAT outputs total copy number (TCN) in contiguous genomic regions, QDNAseq outputs relative copy number (RCN) in 15 Kb bins across the genome and conliga outputs RCN values at specific FAST-SeqS loci. To make a fair comparison between the tools, it was necessary to convert ASCAT’s TCN calls to RCN as follows:

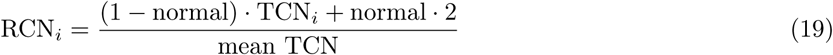

Here, normal represents the estimated normal contamination value provided by ASCAT and *i* represents a contiguous genomic region or a discrete locus or fragment. In the case of a contiguous region, the mean TCN (or ploidy) was calculated as follows:

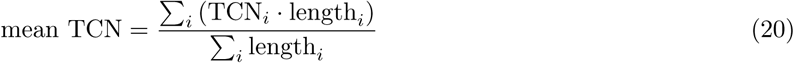

and in the case of discrete loci or fragments:

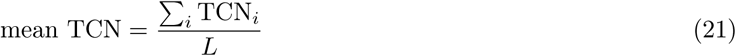

where *L* represents the total number of loci or fragments considered.

In Figure 1e and f, we compared the RCN values at the intersection of genomic loci across ASCAT, QDNAseq and conliga. Since this intersection represented a subset of each method’s genomic loci, the RCN values were rescaled considering only this subset. QDNAseq and conliga RCN values were rescaled by the sample’s mean RCN of the considered loci. ASCAT’s RCN was calculated using equations 19 and 21.

In figure 1g, we compared RCN values in genes of interest. Recurrently amplified and deleted genes were obtained from Dulak *et al.* [23] and Ross-innes *et al.* [12]. Here, ASCAT’s RCN values were calculated using equations 19 and 20 using all called regions for each sample. For each gene in each sample, the weighted mean of the relative copy number (weighted by the length of the overlapping called region) was computed for ASCAT and QDNAseq. This was calculated as follows:

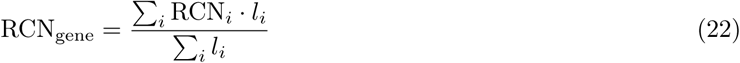

where *l*_*i*_ represents the length of the overlapping portion of the called region with the gene.

For conliga, if loci occurred within the gene, the mean of the RCN values within the gene was used, otherwise the loci directly upstream and downstream, i.e. either side, of the gene were used and a mean value was taken. See Supplementary Table 4 for the full list of genes used in the analysis.

#### Computing Pearson correlation

For each sample, the Pearson correlation coefficient between ASCAT and conliga was calculated. We used ASCAT’s TCN and conliga RCN values at the intersection of genomic loci between ASCAT and conliga. The median value of the sample’s correlation coefficients was reported (all sample correlation coefficients can be found in Supplementary Table 3).

For each sample, the Pearson correlation coefficient between ASCAT and QDNAseq was calculated. We used the intersection of QDNAseq bins with ASCAT copy number regions, using the length-weighted mean of ASCAT’s overlapping TCN values.

When calculating the Pearson correlation for all calls across all samples, we used the re-scaled RCN value at the intersecting genomic loci between ASCAT, QDNAseq and conliga, using the rescaled RCN values described above for Figures 1e and f.

## Code availability

conliga source code is freely available under an open-source GPLv2 license at https://github.com/samabs/ conliga and as Supplementary Software.

## Data availability

The WGS and FAST-SeqS data can be found at the European Genome-phenome Archive (EGA) under accession EGAD00001004289. The copy number results obtained from ASCAT, QDNAseq and conliga can be found https://osf.io/bhx6f/?view_only=ed25e2fb521d46239e5274c032350f0b

## Acknowledgements

Funding for sample sequencing was through the Oesophageal Cancer Clinical and Molecular Stratification (OC-CAMS) Consortium as part of the International Cancer Genome Consortium and was funded by a programme grant from Cancer Research UK (RG66287). SA was funded by Wellcome Trust award [102272]. ERM was funded by MRC Computational Biology Fellowship (MC_UU_12025, MRC Strategic Alliance Funding: MRC Weatherall Institute of Molecular Medicine). We acknowledge the support of The University of Cambridge, Cancer Research UK and Hutchison Whampoa Limited. In particular we acknowledge the support of the Cancer Research UK Cambridge Institute’s Genomics Core Facility. We thank the Human Research Tissue Bank, which is supported by the National Institute for Health Research (NIHR) Cambridge Biomedical Research Centre, from Addenbrooke’s Hospital. Additional infrastructure support was provided from the CRUK funded Experimental Cancer Medicine Centre. The authors also wish to thank Sarah Field, Wing-Kit Leung, Charlie Massie, Paul Coupland and Astrid Wendler for helpful discussions and Ginny Devonshire for her help with uploading the sequencing data to EGA.

## Author contributions statement

RCF, JMJW, SSZ conceived the clinical utility of the experimental approach for upper GI cancer studies. SA, ERM, AGL conceived the broad computational approach for analysis of these experiments. MCVvL, MP, XL, AnN, AJM, JB, AhM, led by SSZ, took the biological samples from patients to data. SA developed and implemented the conliga method, analyzed and preprocessed the data under supervision from ERM and AGL. LB analyzed the high-coverage WGS data. SA wrote the manuscript in consultation with AGL, ERM, RCF and SSZ. All authors read and approved the final version of the manuscript.

## Supplementary Figures

**Supplementary Figure 1.**
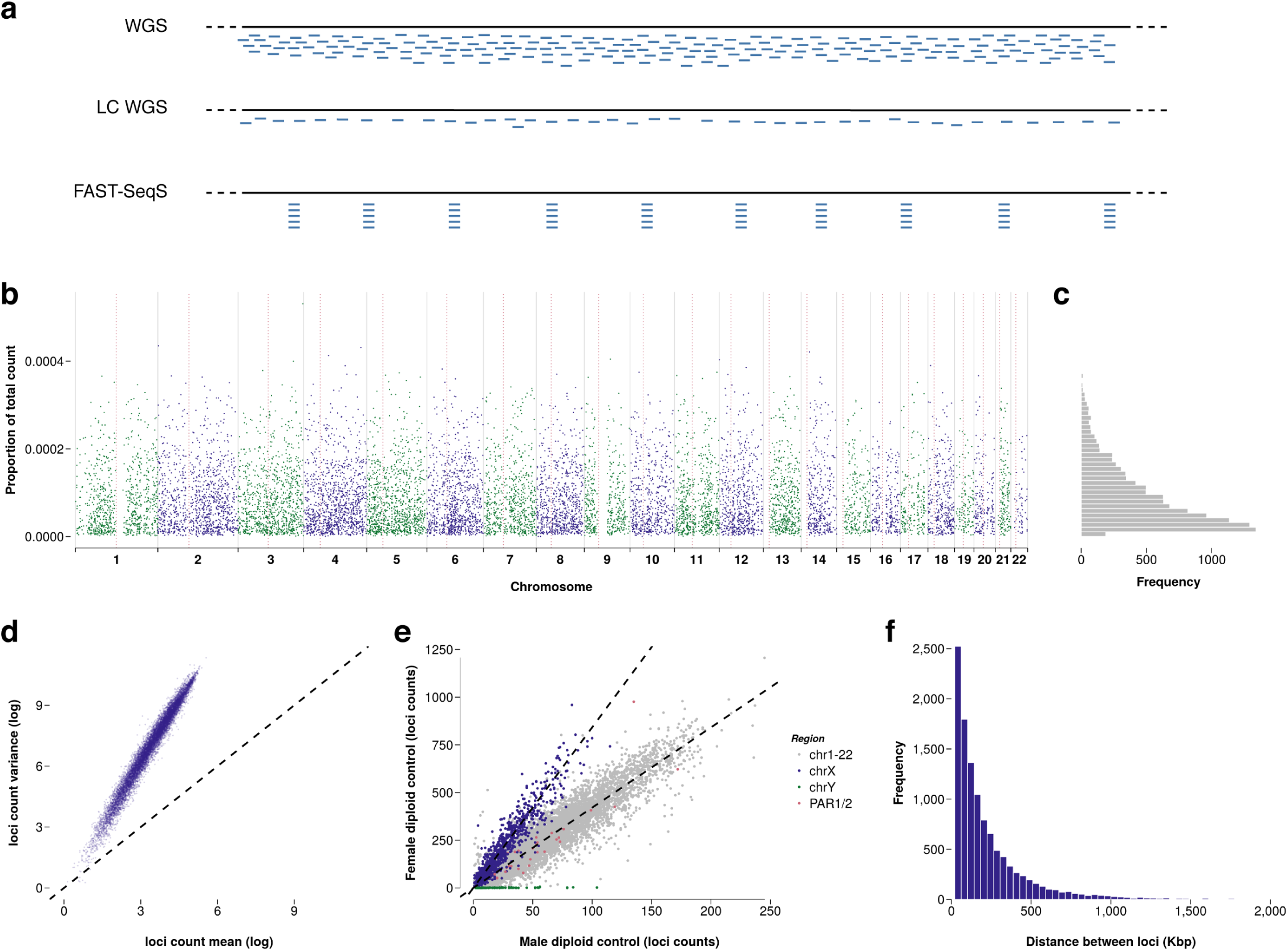
Aspects of FAST-SeqS data. (a) a graphical representation of the different approaches to sequencing for the purposes of SCNA profiling; high-coverage WGS (top), low-coverage WGS (middle), FAST-SeqS (bottom). (b) The proportion of reads obtained at each locus in chr1-22 for control sample (NORM1). (c) Histogram of the proportion of reads obtained at each locus across in chr1-22 for control sample NORM1. (d) log mean vs log variance for each locus in control samples. (e) A male control sample (NORM2) counts plotted against a female control sample (NORM1) counts, showing a relative doubling of count proportions in chrX for the female control sample vs male and absence of counts from chrY in the female sample. (f) Histogram of distances between loci with a mean distance of approximately 200Kbp between loci.

**Supplementary Figure 2.**
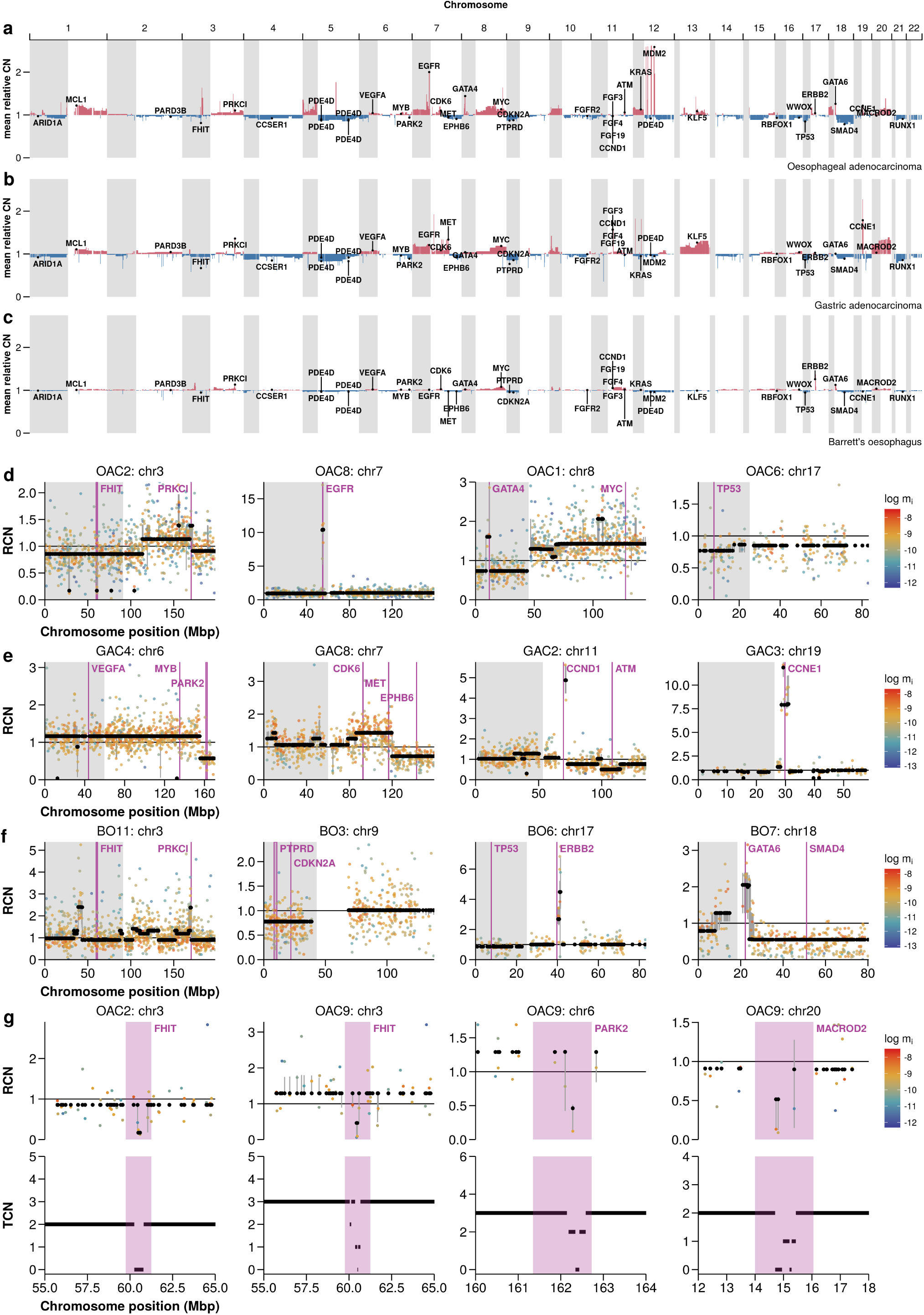
Copy number profile summary of patient cohorts used in this study. (a) Mean relative copy number profile for 11 oesophageal adenocarcinoma samples. (b) Mean relative copy number profile for 8 gastric adenocarcinoma samples. (c) Mean relative copy number profile for 16 Barrett’s oesophagus samples, with varying levels of dysplasia. (d)-(f) Examples of relative copy number profiles for various chromosomes from different samples for OAC, GAC and BO respectively. Black points represent the maximum a posteriori (MAP) relative copy number for each locus, the colored points represent the proportion of reads expected in a control sample (log), with red representing a high proportion and blue representing a low proportion, grey lines represent 90% credible intervals. (g) Zoomed-in regions of chromosomes 3, 6 and 20 showing intra-gene deletion of FHIT, PARK2 and MACROD2. conliga results (top) with comparison to ASCAT (bottom).

